# Sequence to graph alignment using gap-sensitive co-linear chaining

**DOI:** 10.1101/2022.08.29.505691

**Authors:** Ghanshyam Chandra, Chirag Jain

## Abstract

Co-linear chaining is a widely used technique in sequence alignment tools that follow seed-filter-extend methodology. It is a mathematically rigorous approach to combine short exact matches. For colinear chaining between two sequences, efficient subquadratic-time chaining algorithms are well-known for linear, concave and convex gap cost functions [Eppstein *et al*. JACM’92]. However, developing extensions of chaining algorithms for directed acyclic graphs (DAGs) has been challenging. Recently, a new sparse dynamic programming framework was introduced that exploits small path cover of pangenome reference DAGs, and enables efficient chaining [Makinen *et al*. TALG’19, RECOMB’18]. However, the underlying problem formulation did not consider gap cost which makes chaining less effective in practice. To address this, we develop novel problem formulations and optimal chaining algorithms that support a variety of gap cost functions. We demonstrate empirically the ability of our provably-good chaining implementation to align long reads more precisely in comparison to existing aligners. For mapping simulated long reads from human genome to a pangenome DAG of 95 human haplotypes, we achieve 98.7% precision while leaving *<* 2% reads unmapped.

**Implementation:** https://github.com/at-cg/minichain

## 1 Introduction

A significant genetic variation rate among genomes of unrelated humans, plus the growing availability of high-quality human genome assemblies, has accelerated computational efforts to use pangenome reference graphs for common genomic analyses [24,40,41]. The latest version of industry-standard DRAGEN software by Illumina now uses a pangenome graph for mapping reads in highly polymorphic regions of a human genome [14]. For surveys of the recent algorithmic developments in this area, see [2,6,10,33]. Among the many computational tasks associated with pangenome graphs, sequence-to-graph alignment remains a core computational problem. Accurate alignments are required for variation analysis and construction of pangenome graph from multiple genomes [9,22]. Sequence-to-graph alignment is also useful in other applications including genome assembly [13] and long-read error correction [39].

Suppose a pangenome graph is represented as a character labeled DAG *G*(*V, E*) where each vertex *v* ∈ *V* is labeled with a character from alphabet {A,C,G,T}. The sequence-to-DAG alignment problem seeks a path in *G* that spells a string with minimum edit distance from the input query sequence. An *O*(*m* |*E*|) time algorithm for this problem has long been known, where *m* is the length of input sequence [29]. Conditioned on Strong Exponential Time Hypothesis (SETH), the *O*(*m* |*E*|) algorithm is already worst-case optimal up to sub-polynomial improvements because algorithms for computing edit distance in strongly sub-quadratic time cannot exist under SETH [3]. As a result, heuristics must be used for alignment of high-throughput sequencing data against large DAGs to obtain approximate solutions in less time and space.

All practical long read to DAG aligners that scale to large genomes rely on seed–filter–extend methodology [8,22,25,27,34]. The first step is to find a set of *anchors* which indicate short exact matches, e.g., *k*-mer or minimizer matches, between substrings of a sequence to subpaths in a DAG. This is followed by a clustering step that identifies promising subsets of anchors which should be kept within the alignments. Different aligners implement this step in different ways. *Co-linear chaining* is a mathematically rigorous approach to do clustering of anchors. It is well studied for the case of sequence-to-sequence alignment [1,11,12,16,26,28,32], and is widely used in present-day long read to reference sequence aligners [18,21,35,37,38]. For the sequence-to-sequence alignment case, the input to the chaining problem is a set of *N* weighted anchors where each anchor is a pair of intervals in the two sequences that match exactly. A *chain* is defined as an ordered subset of anchors such that their intervals appear in increasing order in both sequences (Figure 1a). The desired output of the co-linear chaining problem is the chain with maximum score where score of a chain is calculated by the sum of weights of the anchors in the chain minus the penalty corresponding to gaps between adjacent anchors. For linear gap costs, this problem is solvable in *O*(*N* log *N*) time by using range-search queries [1].

**Fig. 1:**
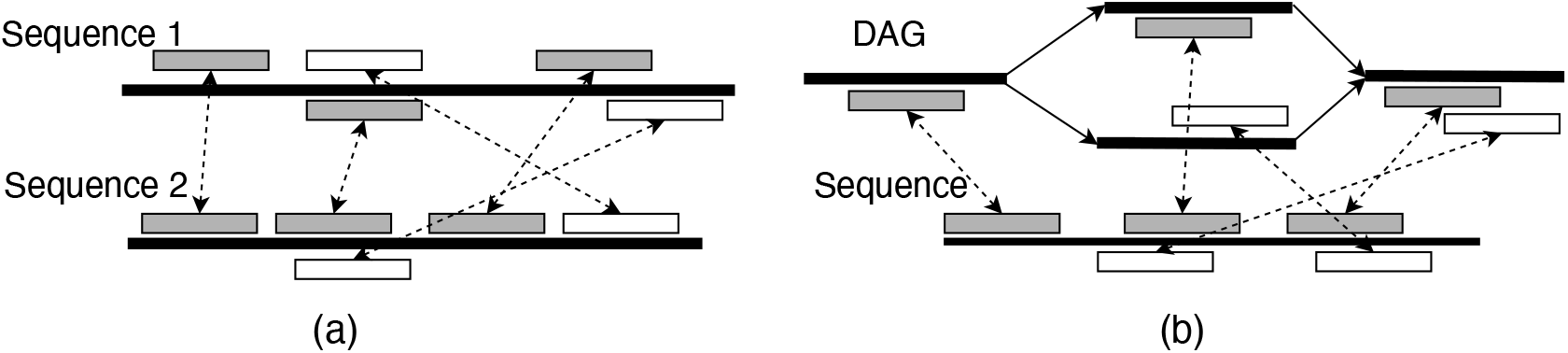
Illustration of co-linear chaining for (a) sequence-to-sequence and (b) sequence-to-DAG alignment. It is assumed that vertices of DAG are labeled with strings. Pairs of rectangles joined by dotted arrows denote anchors (exact matches). A subset of these anchors that form a valid chain are shown in gray.

Solving chaining problem for sequence-to-DAG alignment remained open until Makinen *et al*. [27] introduced a framework that enables sparse dynamic programming on DAGs. Suppose *K* denotes cardinality of a minimum-sized set of paths such that every vertex is covered by at least one path. The algorithm in [27] works by mimicking the sequence-to-sequence chaining algorithm on each path of the *minimum path cover*. After a polynomial-time indexing of the DAG, their algorithm requires *O*(*KN* log *N* + *K* |*V*|) time for chaining. Parameterizing the time complexity in terms of *K* is useful because *K* is expected to be small for pangenome DAGs. This result was further improved in [25] with an *O*(*KN* log *KN*) time algorithm. However, the problem formulations in these works did not include gap cost. Without penalizing gaps, chaining is less effective [16]. A challenge in enforcing gap cost is that measuring gap between two loci in a DAG is not a simple arithmetic operation like in a sequence [20].

We present novel co-linear chaining problem formulations for sequence-to-DAG alignment that penalize gaps, and we develop efficient algorithms to solve them. We carefully design gap cost functions such that they enable us to adapt the sparse dynamic programming framework of Makinen *et al*. [27], and solve the chaining problem optimally in *O*(*KN* log *KN*) time. We implemented and benchmarked one of our proposed algorithms to demonstrate scalability and accuracy gains. Our experiments used human pangenome DAGs built by using 94 high quality *de novo* haplotype assemblies provided by the Human Pangenome Reference Consortium [24] and CHM13 human genome assembly provided by the Telomere-to-Telomere consortium [30]. Using a simulated long read dataset with 0.5× coverage, we demonstrate that our implementation achieves the highest read mapping precision (98.7%) among the existing methods (Minigraph: 98.0%, GraphAligner: 97.0% and GraphChainer: 95.1%). In this experiment, our implementation used 24 minutes and 25 GB RAM with 32 threads, demonstrating that the time and memory requirements are well within practical limits.

## 2 Concepts and notations

### 2.1 Co-linear chaining on sequences revisited

Let *R* and *Q* be two sequences over alphabet *∑* = {*A, C, G, T* }. Let *M*[1..*N*] be an array of anchors. Each anchor is denoted using an interval pair ([*x*..*y*], [*c*..*d*]) with the interpretation that substring *R*[*x*..*y*] matches substring *Q*[*c*..*d*], *x, y, c, d* ∈ ℕ. Anchors are typically either fixed-length matches (e.g., using *k*-mers) or variable-length matches (e.g., maximal exact matches). Suppose function *weight* assigns weights to the anchors. The co-linear chaining problem seeks an ordered subset *S* = *s*_1_*s*_2_ … *s*_*p*_ of anchors from *M* such that

- for all 2 ≤ *j* ≤ *p, s*_*j-*1_ precedes (≺) *s*_*j*_, i.e., *s*_*j-*1_.*y < s*_*j*_.*x*Pand *s*_*j-*1_.*d < s*_*j*_.*c*.
- *S* maximises chaining score, defined as 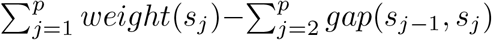. Define *gap*(*s*_*j-*1_, *s*_*j*_) as *f*(*gap*_*R*_(*s*_*j-*1_, *s*_*j*_), *gap*_*Q*_(*s*_*j-*1_, *s*_*j*_)), where *gap*_*R*_(*s*_*j-*1_, *s*_*j*_)= *s*_*j*_.*x* − *s*_*j-*1_.*y* − 1, *gap*_*Q*_(*s*_*j-*1_, *s*_*j*_)= *s*_*j*_.*c* - *s*_*j-*1_.*d* −1 and *f*(*g*_1_, *g*_2_)= *g*_1_ + *g*_2_.

The above problem can be trivially solved in *O*(*N* ^2^) time and *O*(*N*) space. First sort the anchors by the component *M*[·].*x*, and let *T* [1..*N*] be an integer array containing a permutation of set [1..*N*] which specifies the sorted order, i.e.,

*M*[*T* [1]].*x* ≤ *M*[*T* [2]].*x* ≤ … ≤ *M*[*T* [*N*]].*x*. Define array *C*[1..*N*] such that *C*[*j*] is used to store a partial solution, i.e., the score of an optimal chain that ends at anchor *M*[*j*]. Naturally, the final score will be obtained as max_*j*_ *C*[*j*]. Array *C* can be filled by using the following dynamic programming recursion: *C*[*T* [*j*]] = *weight*(*M*[*T* [*j*]]) + max (0, max_*i*:*M*[*i*] ≺*M*[*T* [*j*]]_(*C*[*i*] − *gap*(*M*[*i*],*M*[*T* [*j*]]))), in increasing order of *j*. A straight-forward way of computing *C*[*T* [*j*]] will need an *O*(*N*) linear scan of arrays *C* and *M*, resulting in overall *O*(*N* ^2^) time. However, the *O*(*N* ^2^) algorithm can be optimized to use *O*(*N* log *N*) time by using the following search tree data structure (ref. [4]).

#### Lemma 1.

*Let n be the number of leaves in a balanced binary search tree, each storing a* (*key, value*) *pair. The following operations can be supported in O*(log *n*) *time:*

- *update*(*k, val*): *For the leaf w with key* = *k, value*(*w*) ← max(*value*(*w*), *val*).
- *RMQ*(*l, r*): *Return* max *value*(*w*) | *l < key*(*w*) *< r*}. *This is range maximum query*.

*Moreover, given n* (*key, value*) *pairs, the balanced binary search tree can be constructed in O*(*n* log *n*) *time and O*(*n*) *space*.

The dynamic programming recursion for array *C*[1..*N*] can be computed more efficiently using range maximum queries [1,11]. To achieve this, a search tree needs to be initialized, updated and queried properly (Algorithm 1). Note that argmax_*i*:*M* [*i*] ≺*M* [*j*]_(*C*[*i*] − *gap*(*M*[*i*],*M*[*j*])) is equal to argmax_*i*:*M* [*i*] ≺*M* [*j*]_(*C*[*i*]+ *M*[*i*].*y* + *M*[*i*].*d*). Accordingly, we compute optimal *C*[*j*] in Line 11 by using an *O*(log *N*) time RMQ operation of the form *M*[*i*].*d* ∈ (0,*M*[*j*].*c*) that returns maximum *C*[*i*]+ *M*[*i*].*y* + *M*[*i*].*d* from search tree 𝒯. The algorithm performs *N* update and *N* RMQ operations over search tree 𝒯 of size at most *N*, thus solving the problem in *O*(*N* log *N*) time and *O*(*N*) space.

#### Algorithm 1: *O*(*N* log *N*) time chaining between two sequences

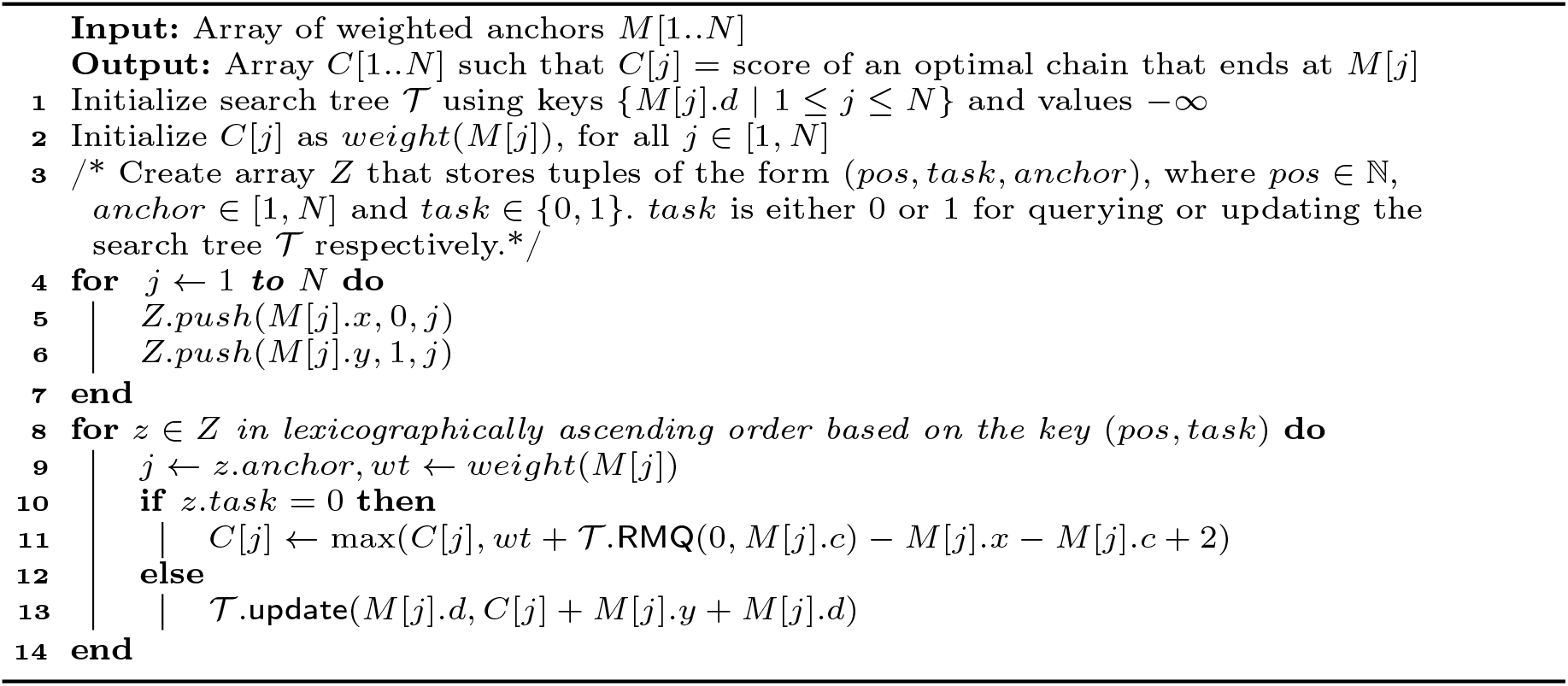

### 2.2 Sparse dynamic programming on DAGs using minimum path cover

Our work builds on the work of Makinen *et al*. [27], who provided a parameterized algorithm to extend co-linear chaining on DAGs without considering gap costs.

In the following, we present useful notations and a summary of their algorithm. In a weighted string-labeled DAG *G*(*V, E, σ*), function *σ* : *V* → *∑*^+^ labels each vertex *v* ∈ *V* with string *σ* (*v*). Edge *v* → *u* has length |*σ* (*v*) |. The length of a path in *G* is the sum of the lengths of the edges traversed in the path. Let *Q* ∈ *∑*^+^ be a query sequence. Let *M* be an array of *N* anchors. An anchor is denoted using a 3-tuple of the form (*v*, [*x*..*y*], [*c*..*d*]) with the interpretation that substring *σ* (*v*)[*x*..*y*] in DAG *G* matches substring *Q*[*c*..*d*], where *x, y, c, d* ∈ ℕ and *v* ∈ *V* (Figure 2). A *path cover* of DAG *G*(*V, E*) is a set of paths in *G* such that every vertex in *V* belongs to at least one path. A *minimum path cover* (MPC) is one having the minimum number of paths. If *K* denotes the size of MPC of DAG *G*, then MPC can be computed either in *O*(*K* |*E*| log |*V*|) [27] or *O*(*K*^3^ |*V*| + |*E*|) [5] time.

**Fig. 2:**
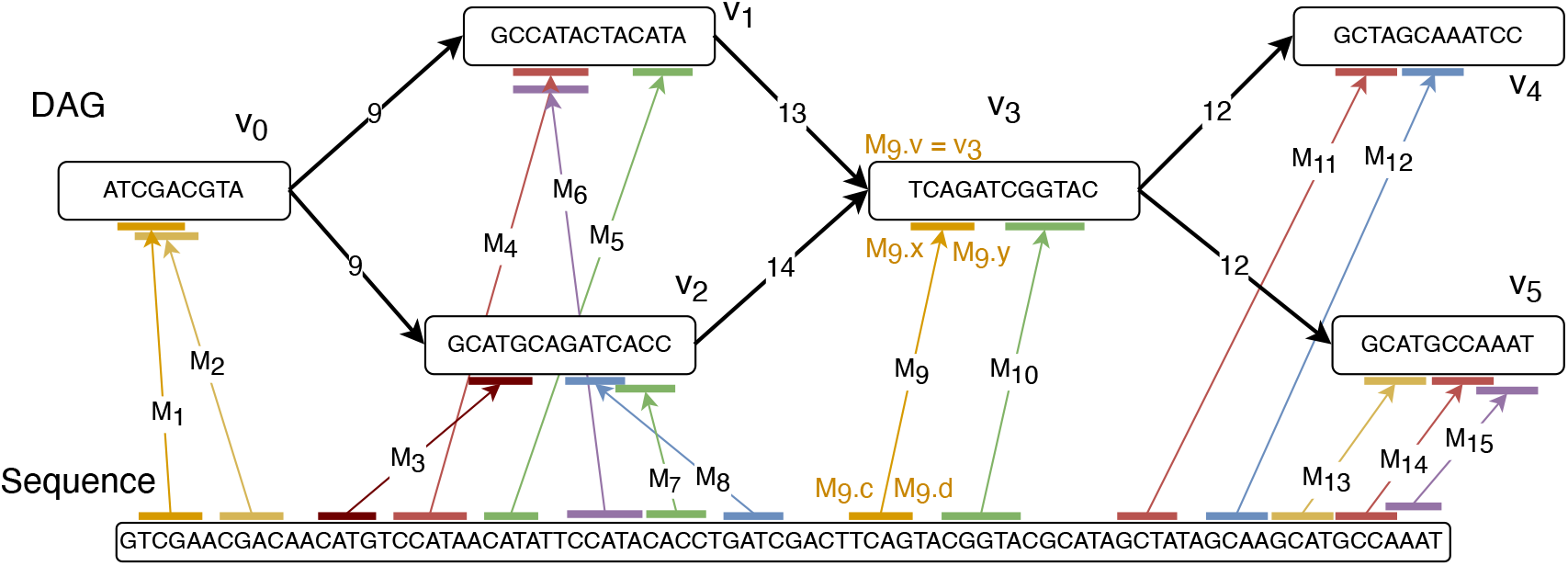
An example showing multiple anchors as input for co-linear chaining. The DAG has a minimum path cover of size two {(*v*_0_, *v*_1_, *v*_3_, *v*_4_), (*v*_0_, *v*_2_, *v*_3_, *v*_5_)}. Anchors *M*_1_, *M*_4_, *M*_5_, *M*_9_, *M*_10_, *M*_11_, *M*_12_ form a valid chain. The interval coordinates of anchor *M*_9_ in the sequence and the DAG are annotated for illustration.

To extend co-linear chaining for sequence-to-DAG alignment, we can use a search tree containing keys equal to the sequence coordinates of anchors, similar to Algorithm 1. However, the order in which the search tree should be queried and updated is not trivial with DAGs. Makinen *et al*. [27] suggested decomposing the DAG into a path cover {*P*_1_, …, *P*_*K*_ }, and then performing the computation only along these paths. The algorithm uses *K* search trees {𝒯_1_, …, 𝒯_*K*_, one per path. Search tree 𝒯_*i*_ maintains *M*[·].*d* as keys and partial solutions *C*[·] as values of all the anchors that lie on path *P*_*i*_. Similar to Algorithm 1, the *K* search trees need to be updated and queried in a proper order. Suppose ℛ (*v*) ⊆ *V* denotes the set of vertices which can reach *v* using a path in *G*. Set ℛ (*v*) always includes *v*. Define *last*2*reach*(*v, i*) as the last vertex on path *P*_*i*_ that belongs to ℛ (*v*), if one exists. Also define *paths*(*v*) as {*i* : *P*_*i*_ covers *v*}. Naturally *last*2*reach*(*v, i*)= *v* iff *i* ∈ *paths*(*v*). The main algorithm works by visiting vertices *u* of *G* in topological order, and executing the following two tasks:

- **Compute optimal scores of all anchors in vertex u**: First, process all the anchors for which *M*[*j*].*v* = *u* in the same order that is used for co-linear chaining on two sequences (Algorithm 1). While performing an update task, update all search trees 𝒯_*i*_, for all *i* ∈ *paths*(*u*). Similarly, while performing a range query, query search trees 𝒯_*i*_ to maximize *C*[*j*].
- **Update partial scores of selected anchors outside vertex u**: Next, for all pairs (*w, i*), *w* ∈ *V, i* ∈ [1, *K*] such that *last*2*reach*(*w, i*) = *u* and *i* ∉ *paths*(*w*), query search tree 𝒯_*i*_ to update score *C*[*j*] of every anchor *M*[*j*] for which *M*[*j*].*v* = *w*.

Based on the above tasks, once the algorithm reaches *v* ∈ *V* in the topological ordering, the scores corresponding to anchors in vertex *v* would have been updated from all other vertices that reach *v*. A well-optimized implementation of this algorithm uses *O*(*KN* log *KN*) time [25]. This result assumes that the DAG is preprocessed, i.e., path cover and *last*2*reach* information is precomputed in *O*(*K*^3^|*V* | + *K*|*E*|) time.

## 3 Problem formulations

We develop six problem formulations for co-linear chaining on DAGs with different gap cost functions. In each problem, we seek an ordered subset *S* = *s*_1_*s*_2_ … *s*_*p*_ of anchors from array *M* such that

- for all 2 ≤ *j* ≤ *p, s*_*j-*1_ precedes (≺) *s*_*j*_, i.e., the following three conditions are satisfied (i) *s*_*j-*1_.*d < s*_*j*_.*c*, (ii) *s*_*j-*1_.*v* ∈ ℛ (*s*_*j*_.*v*), and (iii) *s*_*j-*1_.*y < s*_*j*_.*x* if *s*_*j-*1_.*v* = *s*_*j*_.*v*.
- *S* maximizes the chaining score defined as 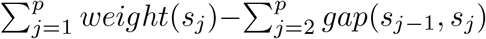. Define *gap*(*s*_*j-*1_, *s*_*j*_) as *f*(*gap*_*G*_(*s*_*j-*1_, *s*_*j*_), *gap*_*S*_(*s*_*j-*1_, *s*_*j*_)), where functions *gap*_*G*_ and *gap*_*S*_ will be used to specify gap cost in the DAG and the query sequence respectively.

*gap*_*S*_(*s*_*j-*1_, *s*_*j*_) equals *s*_*j*_.*c* − *s*_*j-*1_.*d* − 1, i.e., the count of characters in sequence *Q* that occur between the two anchors. However, defining *gap*_*G*_ is not as straightforward because multiple paths may exist from *s*_*j-*1_.*v* to *s*_*j*_.*v*, and the correct alignment path is unknown. We formulate and solve the following problems:

### Problems 1a-1c

*gap*_*G*_(*s*_*j-*1_, *s*_*j*_) is computed by using the shortest path from *s*_*j-*1_.*v* to *s*_*j*_.*v*. Suppose *D*(*v*_1_, *v*_2_) denotes the shortest path length from vertex *v*_1_ to *v*_2_ in *G*. We seek the optimal chaining score when

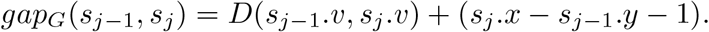

The above expression calculates the count of characters in the string path between anchors *s*_*j-*1_ and *s*_*j*_. Define Problems 1a, 1b and 1c using *f*(*g*_1_, *g*_2_) = *g*_1_ + *g*_2_,*f*(*g*_1_, *g*_2_) = *max*(*g*_1_, *g*_2_) and *f*(*g*_1_, *g*_2_) = |*g*_1_ − *g*_2_| respectively. These definitions of function *f* are motivated from the previous co-linear chaining formulations for sequence-to-sequence alignment [1,28].

### Problems 2a-2c

*gap*_*G*_(*s*_*j-*1_, *s*_*j*_) is measured using a path from *s*_*j-*1_.*v* to *s*_*j*_.*v* that is chosen based on path cover {*P*_1_, …, *P*_*K*_} of DAG *G*. For each *i* ∈ *paths*(*s*_*j-*1_.*v*), consider the following path in *G* that starts from source *s*_*j-*1_.*v* along the edges of path *P*_*i*_ till the middle vertex *last*2*reach*(*s*_*j*_.*v, i*), and finally reaches destination *s*_*j*_.*v* by using the shortest path from *last*2*reach*(*s*_*j*_.*v, i*) to *s*_*j*_.*v*. Among |*paths*(*s*_*j-*1_.*v*)| such possible paths, measure *gap*_*G*_(*s*_*j-*1_, *s*_*j*_) using the path which minimizes *gap*(*s*_*j-*1_, *s*_*j*_)= *f*(*gap*_*G*_(*s*_*j-*1_, *s*_*j*_), *gap*_*S*_(*s*_*j-*1_, *s*_*j*_)). More precisely, *gap*_*G*_(*s*_*j-*1_, *s*_*j*_) equals the element of the set

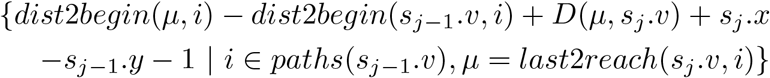

which minimizes *gap*(*s*_*j-*1_, *s*_*j*_), where *dist*2*begin*(*v, i*) denotes the length of sub-path of path *P*_*i*_ from the start of *P*_*i*_ to *v*. We will show that this definition enables significantly faster parameterized algorithms with respect to *K*. Again, define Problems 2a, 2b and 2c with *f*(*g*_1_, *g*_2_)= *g*_1_ + *g*_2_,*f*(*g*_1_, *g*_2_)= *max*(*g*_1_, *g*_2_) and *f*(*g*_1_, *g*_2_)= |*g*_1_ − *g*_2_| respectively.

## 4 Proposed algorithms

Our algorithm to address Problems 1a-1c uses a brute-force approach that evaluates all *O*(*N* ^2^) pairs of anchors. We use single-source shortest distances computations for measuring gaps.

### Lemma 2.

*Problems* 1*a*, 1*b and* 1*c can be solved optimally in O*(*N*(|*V*| + |*E*| + *N*)) *time*.

*Proof*. We will process anchors in array *M*[1..*N*] one by one in a topological order of *M*[·].*v*. If there are two anchors with equal component *M*[·].*v*, then the anchor with lower component *M*[·].*x* is processed first. Suppose DAG *G*′ is obtained by reversing the edges of *G*. While processing anchor *M*[*j*], we will compute partial score *C*[*j*], i.e., the optimal score of a chain that ends at anchor *M*[*j*]. We identify all the anchors that precede *M*[*j*] using a depth-first traversal starting from *M*[*j*].*v* in *G*′. Next, we compute single-source shortest distances from *M*[*j*].*v* in *G*′ which requires *O*(|*V*(| + |*E*|) time for DAGs [7]. Finally, *C*[*j*] is computed as *weight*(*M*[*j*]) + max (0, max_*i*:*M* [*i*] ≺*M* [*j*]_(*C*[*i*] - *f*(*gap*_*G*_(*M*[*i*],*M*[*j*]), *gap*_*S*_(*M*[*i*],*M*[*j*]))) in *O*(*N*) time.

The above algorithm is unlikely to scale to a mammalian dataset. We leave it open whether there exists a faster algorithm to solve Problems 1a-c. Next, we propose *O*(*KN* log *KN*) time algorithm for addressing Problem 2a, assuming *O*(*K*^3^|*V* |+*K*|*E*|) time preprocessing is done for DAG *G*. The preprocessing stage is required to compute (a) an MPC {*P*_1_,…, *P*_*K*_} of *G*, (b) *last*2*reach*(*v, i*), (c) *D*(*last*2*reach*(*v, i*), *v*) and (d) *dist*2*begin*(*v, i*), for all *v* ∈ *V, i* ∈ [1, *K*].

### Lemma 3.

*The preprocessing of DAG G*(*V, E, σ*) *can be done in O*(*K*^3^|*V* | + *K*|*E*|) *time*.

*Proof*. An MPC {*P*_1_,..., *P*_*K*_} can be computed in *O*(*K*^3^ |*V*| + |*E*|) time [5]. To compute the remaining information, we will use dynamic programming algorithms that process vertices ∈ *V* in a fixed topological order. Suppose function *rank* : *V*→ [1, |*V*|] assigns rank to each vertex based on its topological ordering. Let 𝒩 (*v*) denote the set of adjacent in-neighbors of *v*. Similar to [27], *last*2*reach*(*v, i*) is computed in *O*(*K* |*V*| + *K* |*E*|) time for all *v* ∈*V, i* ∈ [1, *K*]. Initialize *last*2*reach*(*v, i*)= 0 for all *v* and *i*. Then, use the following recursion:

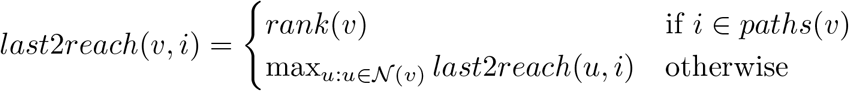

At the end of the algorithm, *last*2*reach*(*v, i*)=0 will hold for only those pairs (*v, i*) for which *last*2*reach*(*v, i*) does not exist. Next, we compute *D*(*last*2*reach*(*v, i*), *v*), for all *v* ∈ *V, i* ∈ [1, *K*], also in *O*(*K*|*V* |+*K*|*E*|) time. Initialize *D*(*last*2*reach*(*v, i*), *v*) = 1 for all *v* and *i*. Then, update *D*(*last*2*reach*(*v, i*), *v*)

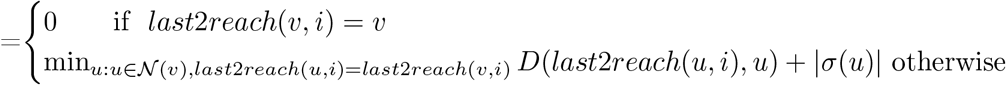

Finally, *dist*2*begin*(*v, i*), for all *v* ∈ *V, i* ∈ [1, *K*] is computed by linearly scanning *K* paths in *O*(*K*|*V* |) time.

### Lemma 4.

*Assuming DAG G*(*V, E, σ*) *is preprocessed, Problem* 2*a can be solved in O*(*KN* log *KN*) *time and O*(*KN*) *space*.

*Proof*. The choice of gap cost definition in Problem 2a allows us to make efficient use of range-search queries. Algorithm 2 gives an outline of the proposed dynamic programming algorithm. Similar to the previously discussed algorithms (Section 2.1), it also saves partial scores in array *C*[1..*N*]. We use *K* search trees, one per path. Search tree 𝒯_*i*_ maintains partial scores *C*[ ] of those anchors *M*[*j*] whose coordinates on DAG are covered by path *P*_*i*_. Each search tree is initialized with keys *M*[*j*].*d*, and values − ∞. Subsequently, *K* search trees are queried and updated in a proper order.

- If *K* = 1, i.e., when DAG *G* is a linear chain, the condition in Line 6 is always satisfied and the term *D*(*v, M*[*j*].*v*) (Line 17) is always zero. In this case, Algorithm 2 works precisely as the co-linear chaining algorithm on two sequences (Algorithm 1).
- For *K >* 1, we use *last*2*reach* information associated with vertex *M*[*j*].*v* (Lines 9-11). This ensures that partial score *C*[*j*] is updated from scores of the preceding anchors on path *P*_*i*_ for all *i* ∈ [1, *K*] \ *paths*(*M*[*j*].*v*).

All the query and update operations done in the search trees together use *O*(*KN* log *N*) time because the count of these operations is bounded by *O*(*KN*), and the size of each tree is ≤*N*. The sorting step in Line 14 requires *O*(*KN* log *KN*) time to sort *O*(*KN*) tuples. The overall space required by *K* search trees and array *Z* is *O*(*KN*).

### Algorithm 2: *O*(*KN* log *KN*) time chaining algorithm for Problem 2a

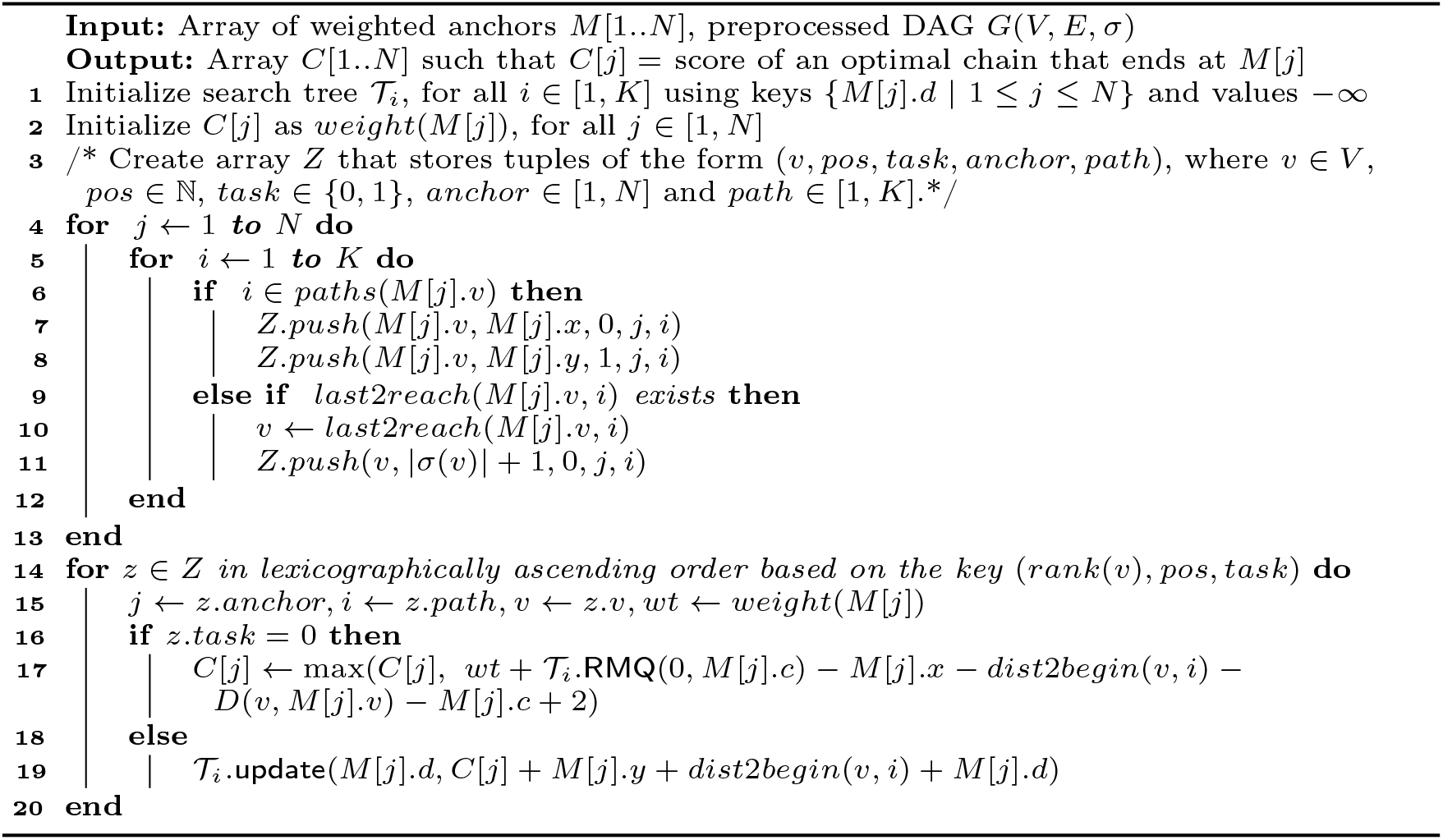

For simplicity of notations, we have not allowed an anchor to span two or more connected vertices in a DAG, but the proposed framework can be easily generalized to handle this [25,27]. Finally, we design algorithms for Problems 2b and 2c by using 2-dimensional RMQs. We summarize the result below and defer the proof to Appendix.

### Lemma 5.

*Assuming DAG G*(*V, E, σ*) *is preprocessed, Problems* 2*b and* 2*c can be solved in O*(*KN* log^2^ *N* + *KN* log *KN*) *time and O*(*KN* log *N*) *space*.

## 5 Implementation details

Among the proposed algorithms, Algorithm 2 has the best time complexity. We implemented this algorithm in `C++`, and developed a practical long read alignment software Minichain.

### Pangenome graph representation

A path in pangenome reference graph *G*(*V, E, σ*) spells a sequence in a single orientation, whereas a read may be sampled from either the same or the opposite orientation due to the double-stranded nature of DNA. To address this internally in Minichain, for each vertex *v* ∈ *V*, we also add another vertex 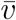 whose string label is the reverse complement of string *σ* (*v*). For each edge *u* → *v* ∈ *E*, we also add the complementary edge 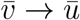. This process doubles the count of edges and vertices.

### Optimization for whole-genome pangenome graphs

A pan-genome reference graph associated with a complete human genome is a union of weakly connected components, one per chromosome, because there is no edge which connects two chromosome components. We actually maintain two components per chromosome, one being the reverse complement of the other. During both preprocessing and chaining stages of the proposed algorithms, each component is treated independently. The parameter *K* in our time complexity results is determined by the maximum *K* value among the components. We use GraphChainer implementation [25] to compute minimum path cover and range queries. We also optimize runtime by performing parallel preprocessing of different components (Lemma 3) using multiple threads.

### Computing multiple best chains and confidence scores

When a read is sampled from a repetitive region of a genome, computing read’s true path of origin becomes challenging. Practical methods often report more than one alignment per read in such cases. The highest-scoring alignment is marked as *primary* alignment, and the remaining are marked as *secondary*. Additionally, based on the score difference between the primary and the highest-scoring secondary alignment, a confidence score ∈ [0, 60] is provided as *mapping quality* that represents the likelihood that the primary alignment is correct [23]. In Minichain, we also implement an algorithm to identify multiple high-scoring chains so that multiple base-to-base alignment records can be reported to a user. Algorithm 2 returns partial scores *C*[1..*N*] in the end. We perform backtracking from anchor argmax_*j*_ *C*[*j*] to compute the optimal chain. The anchors involved in this chain are marked as *visited*. Iteratively, we check presence of another chain (a) whose score is ≥ 𝒯 max_*j*_ *C*[*j*], where 𝒯 ∈ [0, 1] is a user-specified threshold with default value 0.95, and (b) none of the anchors in the chain are previously *visited*. We stop when no additional chains exist that satisfy these conditions.

### Computing anchors and final base-to-base alignments

In Minichain, we use the seeding and base-to-base alignment methods from Minigraph [22]. The seeding method in Minigraph works by identifying common minimizers between query sequence and string labels *σ* (*v*) of graph vertices. Given a pre-defined ordering of all *k*-mers and *w* consecutive *k*-mers in a sequence, (*w, k*)-minimizer is the smallest *k*-mer among the *w k*-mers [36]. The common minimizer occurrences between a query and vertex labels form anchors. In our experiments, we use same parameters *k* = 17,*w* = 11 as Minigraph. The weight of each anchor is *k* times a user-specified constant which is set to 200 by default. Algorithm 2 is used to compute the best chains and discard those anchors which do not contribute to these chains. Finally, we return the filtered anchors to Minigraph’s alignment module to compute base-to-base alignments [42].

## 6 Experiments

### Benchmark datasets

We built string-labeled DAGs of varying sizes by using Minigraph v0.19 [22]. Each DAG is built by using a subset of 95 publicly available haplotype-resolved human genome assemblies [24,30]. In Minigraph, a DAG is iteratively constructed by aligning each haplotype assembly to an intermediate graph, and augmenting additional vertices and/or edges for each structural variant observed. We disabled inversion variants by using `--inv=no` parameter to avoid introducing cycles in the DAG. CHM13 human genome assembly [30] is used as the starting reference, and we added other haplotype assemblies during DAG construction. In the CHM13 assembly, the first 24 contigs represent individual chromosome (1-22, X, Y) sequences, and the last 25^th^ contig represents mitochondrial DNA. Using this data, we constructed five DAGs, labeled as 1H, 10H, 40H, 80H and 95H respectively. In each of these DAG labels, the integer prefix reflects the count of haplotype assemblies present in the DAG. Properties of these DAGs are shown in Table 1. Parameter *K*, i.e., the size of MPC, is presented as a range because different connected components in a DAG have different MPCs. For all DAGs, note that the maximum *K* is ≪ |*V*|. We used PBSIM2 v2.0.1 [31] to simulate long reads from CHM13 human assembly. For each simulated read and each DAG, we know the true string path where the read should align. PBSIM2 input parameters were set such that we get sequencing error rate and N50 read length as 5% and 10 kbp respectively. The commands used to run the different tools are listed in Appendix.

**Table 1:**
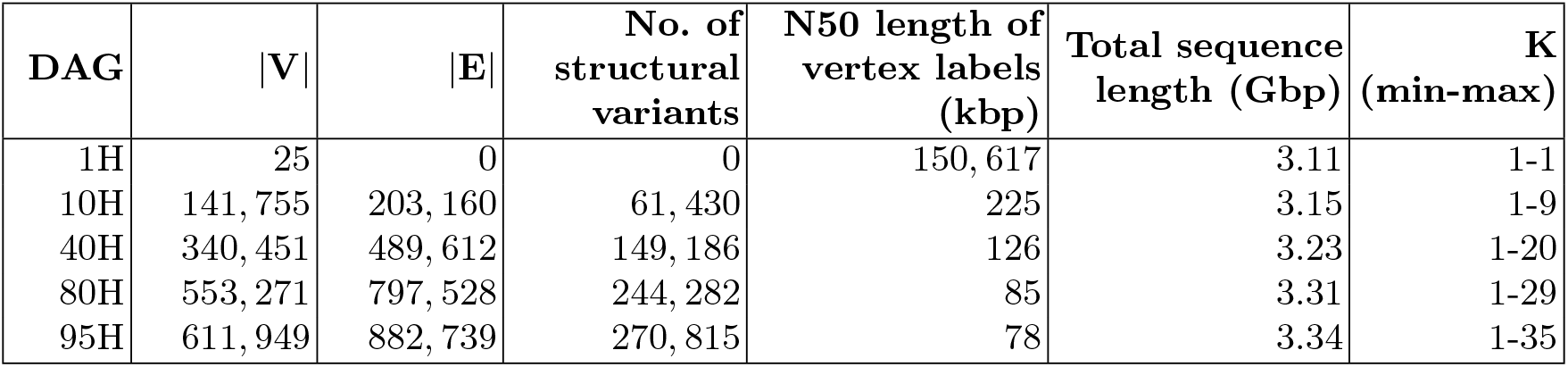
Properties of DAGs used in our experiments. Total sequence length indicates the sum of length of string labels of all vertices in the DAG.

| DAG | $\vert V\vert $ | $\vert E\vert $ | No. of structural variants | N50 length of vertex labels (kbp) | Total sequence length (Gbp) | K (min-max) |
| --- | --- | --- | --- | --- | --- | --- |
| 1H | 25 | 0 | 0 | 150,617 | 3.11 | 1-1 |
| 10H | 141,755 | 203,160 | 61,430 | 225 | 3.15 | 1-9 |
| 40H | 340,451 | 489,612 | 149,186 | 126 | 3.23 | 1-20 |
| 80H | 553,271 | 797,528 | 244,282 | 85 | 3.31 | 1-29 |
| 95H | 611,949 | 882,739 | 270,815 | 78 | 3.34 | 1-35 |

### Evaluation methodology

Alignment output of a read specifies the string path in the input DAG against which the read is aligned. An appropriate evaluation criteria is needed to classify the reported string path as either correct or incorrect by comparing it to the true path. We followed a similar criteria that was used in previous studies [21,22]. First, the reported string path should include only those vertices which correspond to CHM13 assembly, i.e., it should not span an edge augmented from other haplotypes (Figure 3). Second, the reported interval in CHM13 assembly should overlap with the true interval, and the overlapping length should exceed ≥ 10% length of the union of the true and the reported intervals. A correct alignment should satisfy both the conditions. We use `paftools` [21] which implements this evaluation method. All our experiments were done on AMD EPYC 7742 64-core processor with 1 TB RAM. We used 32 threads to run each aligner because all the tested tools support multi-threading by considering each read independently. Wall clock time and peak memory usage were measured using `/usr/bin/time` Linux command.

**Fig. 3:**
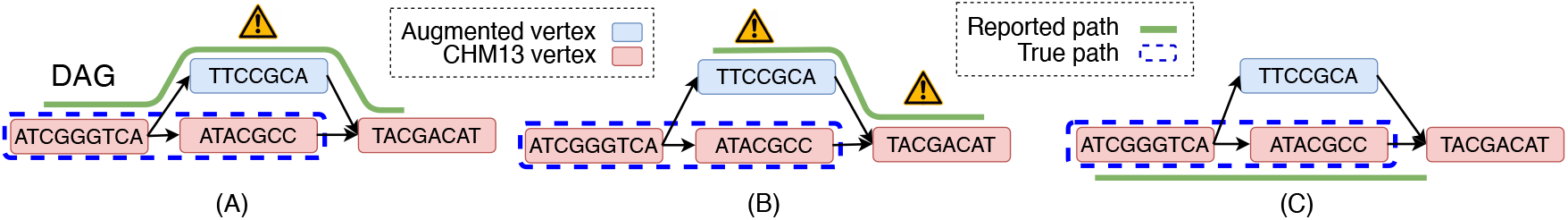
Illustration of the evaluation criteria. Among the three reported paths above, only (C) is correct.

### Performance comparison with existing algorithms

We compared Minichain (v1.0) to three existing sequence-to-DAG aligners: Minigraph v0.19 [22], GraphAligner v1.0.16 [34], and GraphChainer v1.0 [25]. Minigraph uses a two-stage co-linear chaining approach. The first stage ignores edges in the graph and solves colinear chaining between query sequence and vertex labels. The second stage combines the vertex-specific-chains. In contrast, GraphAligner does not use colinear chaining and instead relies on its own clustering heuristics. GraphChainer solves co-linear chaining on DAG without penalizing gaps. All the aligners, except GraphChainer, also compute mapping quality (confidence score) for each alignment. We excluded optimal sequence-to-DAG aligners (e.g., [15,17]) because they do not scale to DAGs built by using entire human genomes.

We evaluated accuracy and runtime of Minichain using three DAGs 1H, 10H and 95H (Tables 2, 3, 4). While using DAG 1H, we also tested Minimap2 v2.24 [21], a well-optimized sequence-to-sequence aligner, by aligning reads directly to CHM13 genome assembly. The results show that Minichain consistently achieves the highest precision among the existing sequence-to-DAG aligners. It aligns a higher fraction of reads compared to Minigraph. The gains are also visible when mapping quality (MQ) cutoff 10 is used to filter out low-confidence alignments. GraphAligner and GraphChainer align 100% reads consistently, but this is supplemented with much higher fraction of incorrectly aligned reads. Both Minigraph and Minichain do not align 100% reads. This likely happens because the seeding method used in these two aligners filters out the most frequently occurring minimizers from DAG to avoid processing too many anchors. This can leave several reads originating from long-repetitive genomic regions as unaligned [19].

**Table 2:** Performance comparison of long read aligners using DAG 1H.

|  | Minichain | Minigraph | GraphAligner | GraphChainer | Minimap2 |
| --- | --- | --- | --- | --- | --- |
| Indexing time (sec) | **65** | **65** | 395 | 458 | 55 |
| Alignment time (sec) | 205 | **104** | 6298 | 6714 | 133 |
| Memory usage (GB) | 20.06 | **19.66** | 38.12 | 126.14 | 12.48 |
| Unaligned reads | 0.94% | 2.11% | **0%** | **0%** | 0% |
| Incorrect aligned reads | **0.58%** | 0.66% | 1.06% | 1.33% | 0.56% |
| Unaligned reads (MQ≥10) | 3.89% | 5.82% | 0.80% | **0%** | 2.29% |
| Incorrect aligned reads (MQ≥10) | **0.02%** | 0.11% | 0.53% | 1.33% | 0.0013% |

**Table 3:** Performance comparison of long read aligners using DAG 10H.

|  | Minichain | Minigraph | GraphAligner | GraphChainer |
| --- | --- | --- | --- | --- |
| Indexing time (sec) | 67 | **66** | 321 | 537 |
| Alignment time (sec) | 610 | **132** | 5479 | 9642 |
| Memory usage (GB) | **23.15** | 23.16 | 38.41 | 143.94 |
| Unaligned reads | 1.17% | 2.17% | **0%** | **0%** |
| Incorrect aligned reads | **0.80%** | 1.20% | 1.55% | 2.10% |
| Unaligned reads (MQ≥10) | 4.03% | 5.88% | 0.28% | **0%** |
| Incorrect aligned reads (MQ≥10) | **0.20%** | 0.34% | 0.99% | 2.10% |

**Table 4:** Performance comparison of long read aligners using DAG 95H.

|  | Minichain | Minigraph | GraphAligner | GraphChainer |
| --- | --- | --- | --- | --- |
| Indexing time (sec) | 77 | **71** | 342 | 763 |
| Alignment time (sec) | 1414 | **154** | 5695 | 17336 |
| Memory usage (GB) | **24.75** | 24.76 | 40.79 | 192.36 |
| Unaligned reads | 1.62% | 2.23% | **0%** | **0%** |
| Incorrect aligned reads | **1.31%** | 1.96% | 3.01% | 4.92% |
| Unaligned reads (MQ≥10) | 4.75% | 6.26% | 0.88% | **0%** |
| Incorrect aligned reads (MQ≥10) | **0.56%** | 0.89% | 2.38% | 4.92% |

Among the four aligners, Minigraph performs the best in terms of runtime. Runtime of Minichain increases for DAG 95H because of higher value of *K*. However, we expect that this can be partly addressed with additional improvements in the proposed chaining algorithm, e.g., by dynamically deleting the anchors from search trees whose gap from all the remaining unprocessed anchors exceeds an acceptable limit. Overall, the results demonstrate practical advantage of Minichain for accurate long-read alignment to DAGs. Superior accuracy of Minichain is also illustrated using precision-recall plots in Figure 4.

**Fig. 4:**
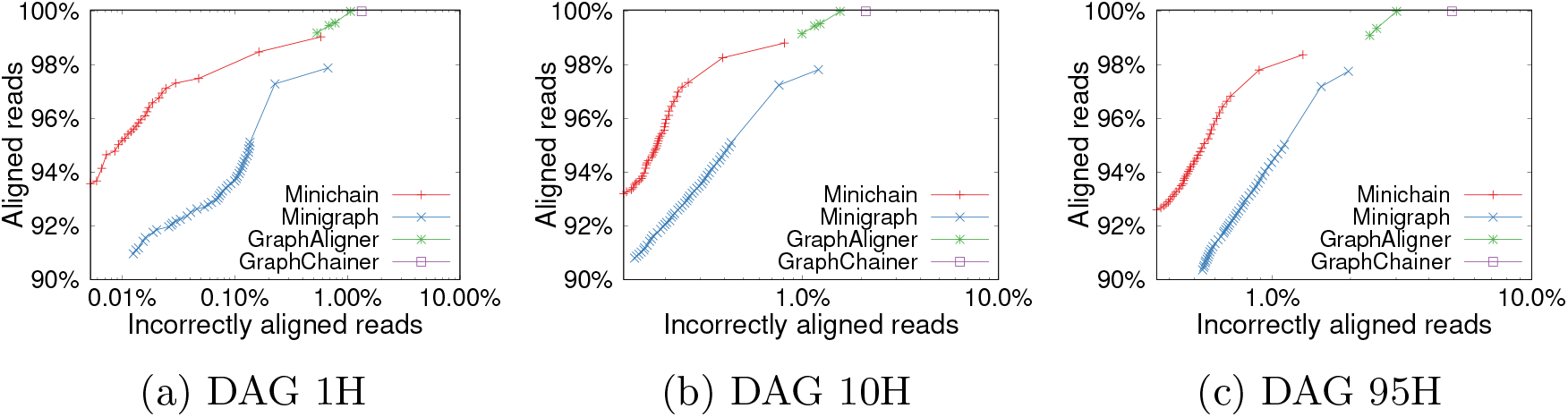
Precision-recall curves obtained by using different aligners. X-axis indicates percentage of incorrectly aligned reads in log-scale. These curves are obtained by setting different mapping quality cutoffs ∈ [0, 60]. GraphChainer curve is a single point because it reports fixed mapping quality 60 in all alignments.

### Impact of increasing DAG size on accuracy

Alignment accuracy generally deteriorates as count of haplotypes increases in DAGs for all the tested aligners. For each read that was not aligned correctly, we checked if the corresponding reported string path overlaps with the true interval (Figure 3, case A). Such reads are aligned to *correct region* in the DAG but the reported path uses one or more augmented edges. The remaining fraction of incorrectly aligned reads align to *wrong region* in the DAG. We observe that the fraction of incorrectly-aligned reads which align to correct region in DAG increases with increasing count of haplotypes (Figure 5). This happens because the count of alternate paths increases combinatorially with more number of haplotypes which makes precise alignment of a read to its path of origin a challenging problem. Addressing this issue requires further algorithmic improvements.

**Fig. 5:**
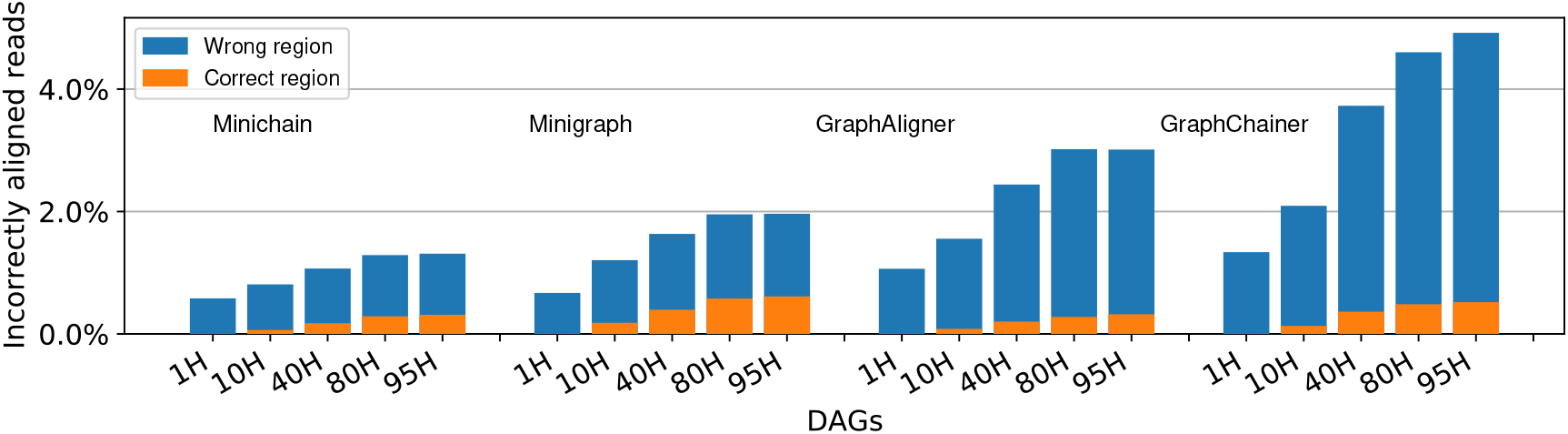
The fraction of incorrectly aligned reads is shown using DAGs 1H, 10H, 40H, 80H and 95H. Each incorrectly-aligned read is further classified as aligned to either a wrong or a correct region in the DAG based on whether the reported string path overlaps with the true string path (e.g., cases A,B in Figure 3).

## Acknowledgements

This work was supported by funding from the National Supercomputing Mission, India under DST/NSM/ R&D_HPC_Applications. We used computing resources provided by the C-DAC National PARAM Supercomputing Facility, India, and the National Energy Research Scientific Computing Center, USA.

## Appendix

### 1 Algorithms for solving Problems 2b and 2c

To support Lemma 5, we propose *O*(*KN* log^2^ *N* + *KN* log *KN*) time algorithms to solve Problems 2b and 2c. We assume that DAG *G*(*V, E, σ*) is preprocessed already (Lemma 3). Unlike Algorithm 2 which uses 1D orthogonal range queries, here we will use 2D orthogonal range queries. When function *f*(*gap*_*G*_(*s*_*j-*1_, *s*_*j*_), *gap*_*S*_(*s*_*j-*1_, *s*_*j*_)) for gap cost is defined as either max(*gap*_*G*_(*s*_*j-*1_, *s*_*j*_), *gap*_*S*_(*s*_*j-*1_, *s*_*j*_)) or |*gap*_*G*_(*s*_*j-*1_, *s*_*j*_) − *gap*_*S*_(*s*_*j-*1_, *s*_*j*_)|, the second dimension is used to check whether *gap*_*G*_(*s*_*j-*1_, *s*_*j*_) ≥ *gap*_*S*_(*s*_*j-*1_, *s*_*j*_) holds or not. The following search tree data structure is used to support 2D orthogonal range queries (ref. [4]).

#### Lemma 6.

*Let n be the number of* (*key, value*) *entries where a key is defined as a tuple* (*k*_1_, *k*_2_) *of type* ℤ × ℤ. *The following operations can be supported in O*(log^2^ *n*) *time:*

- *update*((*k*_1_, *k*_2_), *val*): *For the entry w with key* = (*k*_1_, *k*_2_), *value*(*w*) ← max(*value*(*w*), *val*).
- *RMQ*((*l*_1_, *r*_1_), (*l*_2_, *r*_2_)): *Return* max {*value*(*w*) | *l*_1_ *< key*(*w*).*k*_1_ *< r*_1_, *l*_2_ *< key*(*w*).*k*_2_ *< r*_2_ }. *This is* 2*-dimensional range maximum query. Input ranges can be provided as either open or closed intervals*.

*Given n* (*key, value*) *entries, a search tree data structure to support the above operations can be constructed in O*(*n* log *n*) *time and space*.

Algorithm 3 outlines our solution to Problem 2b. The pseudocode follows a similar structure as Algorithm 2. We use 2*K* search trees 𝒯_*i*_ and ℐ_*i*_, for all *i* ∈ [1, *K*] (Line 1). Search trees 𝒯_*i*_ execute update operations to handle those cases where gap cost should equal *gap*_*S*_(*s*_*j-*1_, *s*_*j*_) (Line 13). In Line 8, the first dimension of the range query in 𝒯_*i*_ is to restrict search over those anchors which precede anchor *M*[*j*] (same as Algorithm 2), and the second dimension is to further restrict search over those preceding anchors *M*[*i*] for which *gap*_*S*_(*M*[*i*],*M*[*j*]) ≥ *gap*_*G*_(*M*[*i*],*M*[*j*]). Similarly, search trees ℐ_*i*_ set value of each key to handle cases where gap cost equals *gap*_*G*_(*s*_*j-*1_, *s*_*j*_) (Line 14). In Line 9, the first dimension of the range query in ℐ_*i*_ is to restrict search over those anchors which precede anchor *M*[*j*], and the second dimension further restricts search over those anchors *M*[*i*] for which *gap*_*S*_(*M*[*i*],*M*[*j*]) ≤ *gap*_*G*_(*M*[*i*],*M*[*j*]). Throughout the algorithm, we perform *O*(*KN*) RMQ and update operations. Each operation uses *O*(*log*^2^*N*) time because size of each tree is at most *N*. Therefore, these operations require overall *O*(*KN* log^2^ *N*) time. The sorting operation used in Line 4 uses *O*(*KN* log *KN*) time to sort *O*(*KN*) tuples. The combined storage complexity of 2*K* search trees is *O*(*KN* log *N*). This completes the description of Algorithm 3. Algorithm 4 addresses Problem 2c, and follows a similar intuition.

#### Algorithm 3: *O*(*KN* log^2^ *N* + *KN* log *KN*) time co-linear chaining algorithm to solve Problem 2b

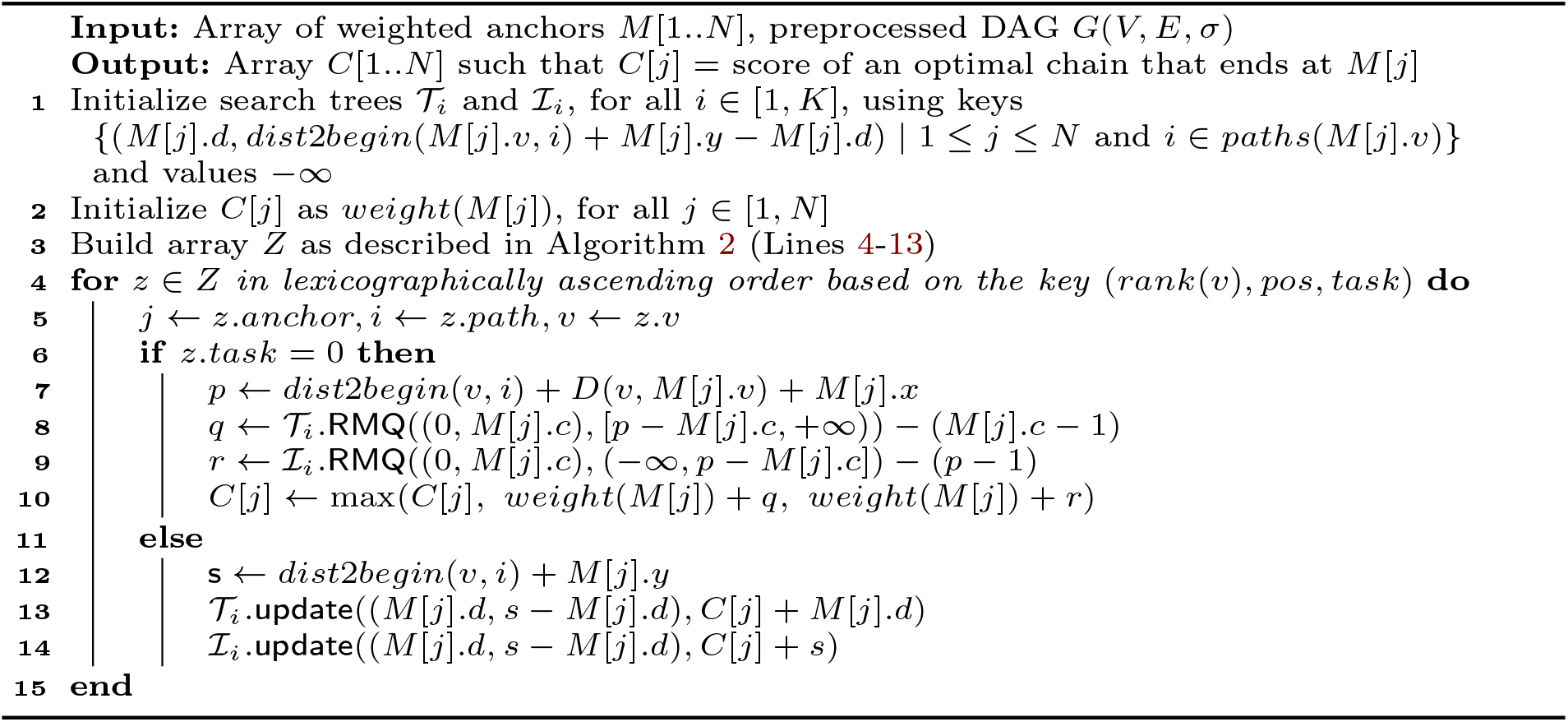

#### Algorithm 4: *O*(*KN* log^2^ *N* + *KN* log *KN*) time co-linear chaining algorithm to solve Problem 2c

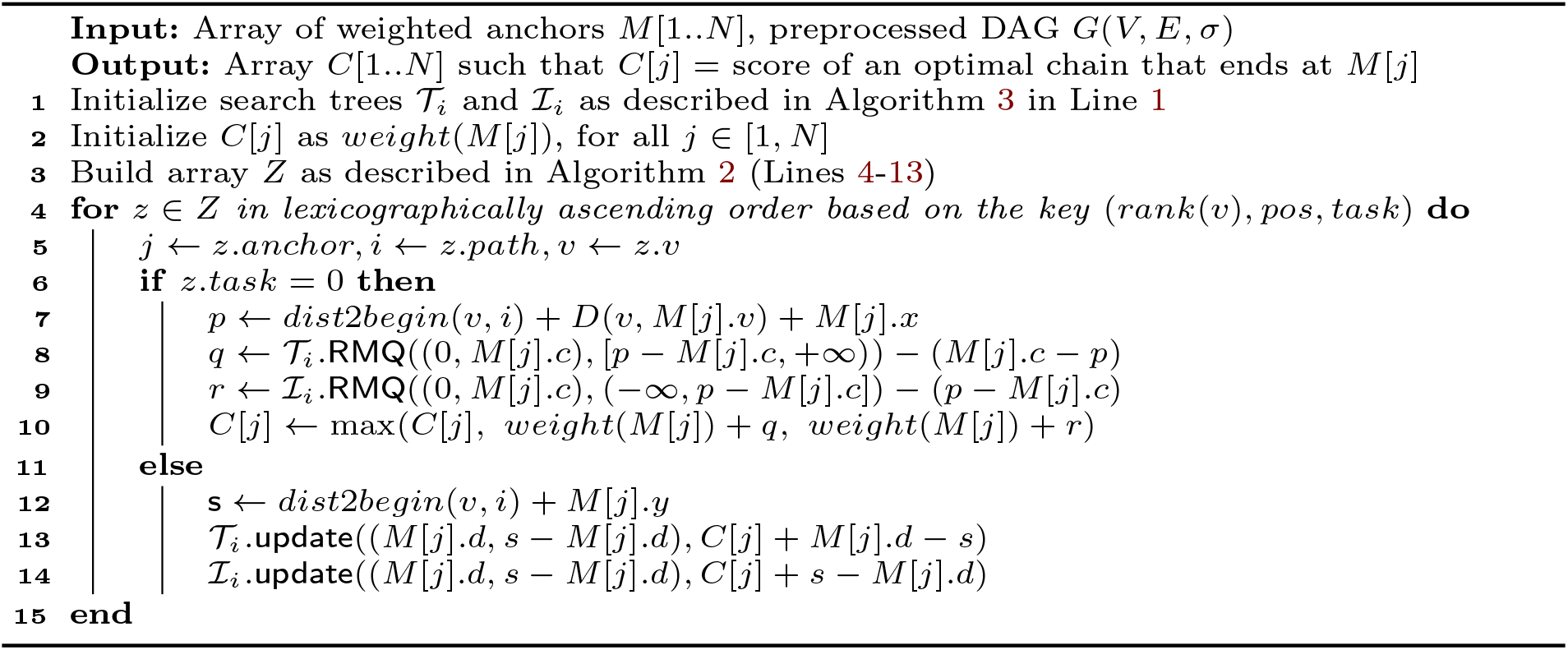

### 2 Commands used for empirical evaluation

**Table 5:** Command-line arguments used to run various tools. The scripts to reproduce results are available in Minichain GitHub repository.

| Purpose | Tool | Commands |
| --- | --- | --- |
| Read simulation | pbsim2 (v2.0.1, bioconda)<br>paftools.js (commit:15cade0) | pbsim --depth 0.5 --length-min 10000 --length-mean 10000 --length-max 10000 --accuracy-mean 0.95 --accuracy-max 0.95 --accuracy-min 0.95 --hmm_model P6C4.model CHM13Y.fa<br>k8-Linux paftools.js pbsim2fq CHM13Y.fa.fai *.maf > CHM13Y_reads.fa |
| Graph generation | minigraph (v0.19-r551) | minigraph -t32 -cxggs --inv=no CHM13Y.fa HG002.1.fa HG002.2.fa HG00438.1.fa HG00438.2.fa HG005.1.fa HG005.2.fa HG00621.1.fa HG00621.2.fa HG00673.1.fa > CHM13Y_10H.gfa 2> log_10H.txt |
| Read mapping | minimap2 (v2.24-r1122)<br>minigraph (v0.19-r551)<br>minichain (v1.0)<br>GraphAligner (commit:4e2ca66)<br>GraphChainer (commit:59c9c67) | minimap2 -t32 -cx map-pb CHM13Y.fa CHM13Y_reads.fa > CHM13Y.paf<br>minigraph -t32 -cx lr CHM13Y_10H.gfa CHM13Y_reads.fa > CHM13Y_10H.gaf<br>minichain -t32 -cx lr CHM13Y_10H.gfa CHM13Y_reads.fa > CHM13Y_10H.gaf<br>GraphAligner -t32 -x vg -g CHM13Y_10H.gfa -f CHM13Y_reads.fa -a CHM13Y_10H.gaf<br>GraphChainer -t32 -g CHM13Y_10H.gfa -f CHM13Y_reads.fa -a CHM13Y_10H.gaf |
| Conversion to stable GAF co-ordinates (GraphAligner and GraphChainer) | mgutils.js (commit:8619249) | k8-Linux mgutils.js stableGaf CHM13Y_10H.gfa CHM13Y_10H.gaf > CHM13Y_10H_stable.gaf |
| Evaluation of read alignments | paftools.js | k8-Linux paftools.js mapeval CHM13Y_10H.gaf |

## Notes

### Competing Interest Statement

The authors have declared no competing interest.

### Summary of Updates

Revised the Naive Algorithm.

